# FlyWire: Online community for whole-brain connectomics

**DOI:** 10.1101/2020.08.30.274225

**Authors:** Sven Dorkenwald, Claire McKellar, Thomas Macrina, Nico Kemnitz, Kisuk Lee, Ran Lu, Jingpeng Wu, Sergiy Popovych, Eric Mitchell, Barak Nehoran, Zhen Jia, J. Alexander Bae, Shang Mu, Dodam Ih, Manuel Castro, Oluwaseun Ogedengbe, Akhilesh Halageri, Zoe Ashwood, Jonathan Zung, Derrick Brittain, Forrest Collman, Casey Schneider-Mizell, Chris Jordan, William Silversmith, Christa Baker, David Deutsch, Lucas Encarnacion-Rivera, Sandeep Kumar, Austin Burke, Jay Gager, James Hebditch, Selden Koolman, Merlin Moore, Sarah Morejohn, Ben Silverman, Kyle Willie, Ryan Willie, Szi-chieh Yu, Mala Murthy, H. Sebastian Seung

## Abstract

Due to advances in automated image acquisition and analysis, new whole-brain connectomes beyond *C. elegans* are finally on the horizon. Proofreading of whole-brain automated reconstructions will require many person-years of effort, due to the huge volumes of data involved. Here we present FlyWire, an online community for proofreading neural circuits in a fly brain, and explain how its computational and social structures are organized to scale up to whole-brain connectomics. Browser-based 3D interactive segmentation by collaborative editing of a spatially chunked supervoxel graph makes it possible to distribute proofreading to individuals located virtually anywhere in the world. Information in the edit history is programmatically accessible for a variety of uses such as estimating proofreading accuracy or building incentive systems. An open community accelerates proofreading by recruiting more participants, and accelerates scientific discovery by requiring information sharing. We demonstrate how FlyWire enables circuit analysis by reconstructing and analysing the connectome of mechanosensory neurons.

## INTRODUCTION

Electron microscopy (EM) is the only technique that so far has proven capable of reconstructing all connections in a nervous system or brain. While the activity of large populations of neurons or even entire vertebrate brains (Ahrens et al. 2013) can be observed via calcium imaging, adult connectomes have so far been mapped for only one species, *C. elegans* (White et al. 1986; Cook et al. 2019). Fortunately, connectomes of more complex brains are now finally on the horizon. A major milestone in this direction has been the recent release of a *Drosophila* “hemibrain” connectome (Xu et al. 2020). Part of a fly brain was imaged by EM, and automatically reconstructed using deep learning. Many errors in the reconstruction were corrected by 50 person-years of human “proofreading” to create a first draft of the hemibrain connectome.

The entire fly brain connectome would be of great scientific interest, because *Drosophila melanogaster* has become a popular model organism for circuit neuroscience. Flies are capable of a wide array of complex behaviors and functions, including social communication, aggression, spatial navigation, decision-making, and learning (Coen et al. 2014; Duistermars et al. 2018; Seelig and Jayaraman 2015; DasGupta, Ferreira, and Miesenböck 2014; Owald et al. 2015). While the hemibrain connectome is proving extremely useful for *Drosophila* circuit neuroscience, circuits that extend outside the hemibrain volume cannot be reconstructed (Extended Figure 1).

Therefore, we have created FlyWire, the first open online community for proofreading a connectome of a whole brain (flywire.ai). FlyWire is based on an EM dataset of a full adult fly brain (FAFB) that was previously released (Zheng et al. 2018). While FlyWire is dedicated to the fly brain, it introduces several new methods that will be generally applicable to whole-brain connectomics. The first is a novel data structure called the ChunkedGraph, which is the basis for proofreading. Like previous systems (J. S. Kim et al. 2014; Haehn et al. 2014; Knowles-Barley et al. 2016; Zhao et al. 2018), FlyWire represents neurons as connected components in a graph of supervoxels (groups of voxels). A naive implementation of this underlying data structure would scale poorly to large datasets. The ChunkedGraph divides the graph into spatially local chunks, and adds a hierarchy of extra vertices and edges to cache information about connected components. We show that edit operations are over an order of magnitude faster than systems relying on the naive implementation of the supervoxel graph. In addition, the ChunkedGraph enables real-time collaboration and stores the history of all edits.

FlyWire is also novel for its open social structure. Membership in the community is open to everyone. Community members immediately share the results of proofreading with each other. In contrast, members of the “walled garden” community, also reconstructing circuits from the FAFB dataset (Felsenberg et al. 2018; Dolan et al. 2018), are selected to avoid conflicts between labs working on the same circuit. Rather than restrict membership, FlyWire attempts to avoid conflicts by enforcing sharing of reconstructions with attribution. The hemibrain was reconstructed through a closed proofreading process that mobilized paid workers, and updated results are released to the public as the internal proofreading progresses (Xu et al. 2020). Our principle of openness was inspired by a previous project to reconstruct larval *Drosophila* circuits (A. Cardona, personal communication).

The walled garden has historically used fully manual skeletonization to reconstruct neural circuits from FAFB (Felsenberg et al. 2018; Dolan et al. 2018). Since manual skeletonization is laborious, the walled garden is starting to migrate to semi-automated reconstruction (Z. Zheng et al. 2020) based on piecing together automatically generated skeletons (Li et al. 2019). FlyWire, in addition to being open, enables true 3D interactive proofreading of a volumetric segmentation.

A final novelty of FlyWire is that the accuracy of the automated reconstruction was boosted by realigning the serial section images using deep learning (Mitchell et al. 2019). In the published FAFB dataset, which was aligned with conventional computer vision algorithms (Zheng et al. 2018), misalignments were numerous enough to be the dominant failure mode for automated reconstruction.

We show that most neurons in FlyWire can be reconstructed within 1.5 hours to a degree that is sufficient for morphological and circuit level analyses. Using FlyWire we produced the first complete connectivity diagram between known early mechanosensory/auditory neurons and discovered previously unknown connection patterns, including cross-hemispheric connections likely to be important for sound localization. As further evidence of FlyWire’s utility, it was also recently used to map the connectivity of *Drosophila* neurons related to a persistent internal state (Deutsch, Pacheco, and Encarnacion-Rivera 2020).

Given recent progress in automated EM image acquisition (Yin et al. 2019) reconstruction (Jain, Seung, and Turaga 2010; Lee et al. 2019; Januszewski et al. 2018), we can expect many other large scale connectome projects in the future. The methods introduced by FlyWire should be generally useful for these projects.

## RESULTS

### Neuron segmentation

We realigned the serial section images of the FAFB dataset (Zheng et al. 2018), and generated an automated segmentation (see Methods, Extended Data Fig. 2 for a quality assessment of the realignment). The automatically generated segments often show many or all of the expected parts of a fly neuron: a soma, dendrites, axon terminals, and a primary neurite (the usually unbranched proximal neurite connecting the soma to branching arbors downstream).

We examined reconstructions of well known cell types before and after proofreading to assess segmentation quality and test whether neurons can be identified before proofreading (Fig. 1). Indeed, the automated segmentation is often highly accurate to begin with (quantification below) and unique morphological features across the examined cells are visible without proofreading. Qualitative comparison between images of light-level stains of the giant fiber neurons (Pézier et al. 2016) (Fig. 1 a,c,e) and a mushroom body APL neuron (Wu et al. 2011) (Fig. 1 b,d,f) show that our semi-automated segmentation procedures are able to capture neurons in their entirety.

**Figure 1.**
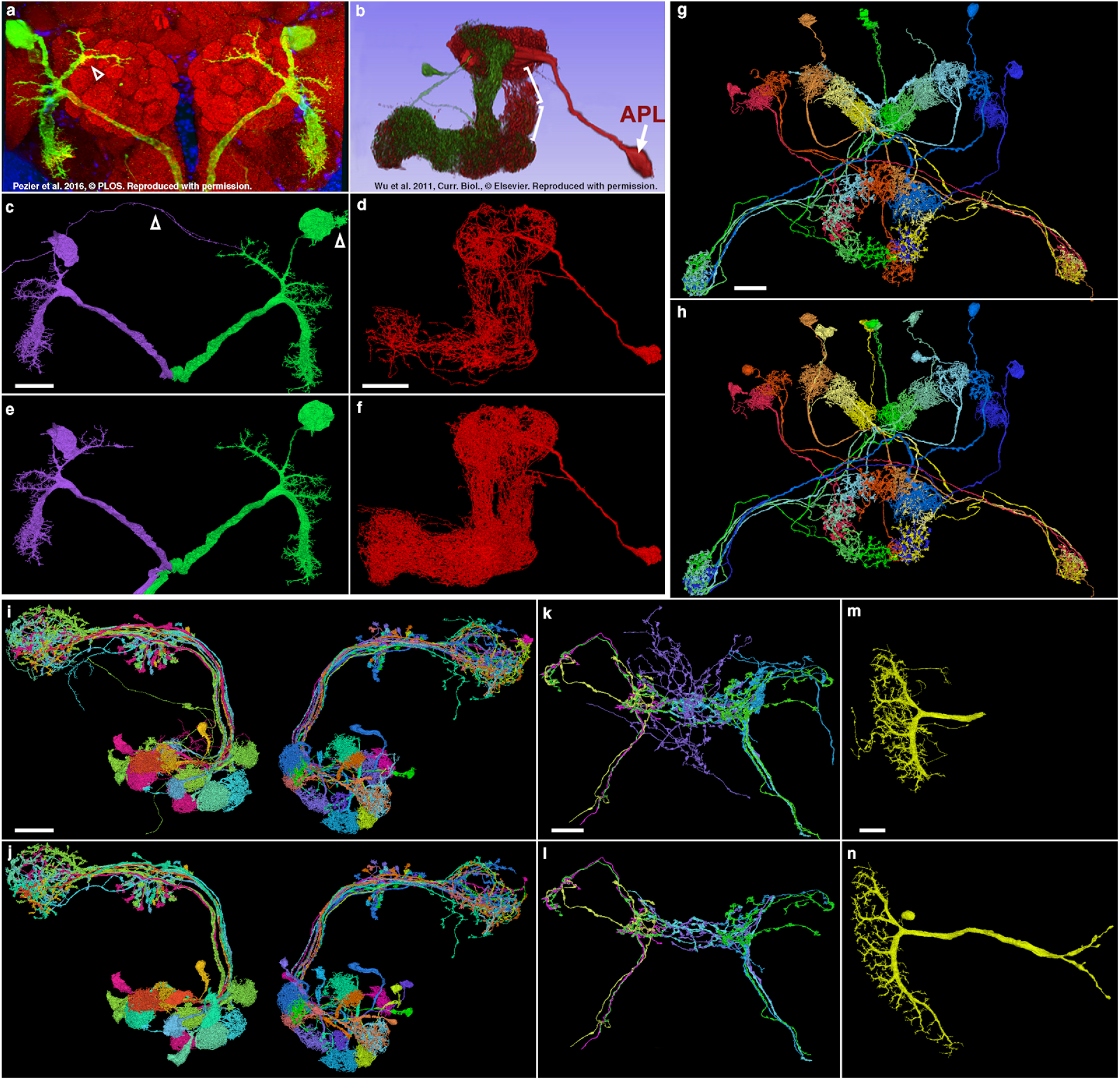
Assessing segmentation quality using known neurons. (**a-d**) Comparison of light-level stains of known giant fiber neurons (**a**) and a mushroom body APL neuron (b, red) to FlyWire’s AI-predicted segmentation of these cells (**c,d**). (**e,f**) The same neurons shown following proofreading. (**g-n**) Examples of other known cell types before and after proofreading (top and bottom of each pair, respectively): central complex neurons (g,h), olfactory projection neurons (**i,j**), gustatory receptor neurons (**k,l**) and a lobula plate tangential cell (**m,n**). All views frontal except APL and central complex neurons: dorso-frontal view. Scale bars: (c, d, e, f, i, j) 30 μm, (g, h, k, l) 15 μm, (m, n) 20 μm

### Chunked supervoxel graph as data structure for proofreading

Proofreading consists of two basic operations: merging falsely disconnected segments and splitting falsely merged ones. For efficient editing of the automatically generated segments, we represent the segmentation as a supervoxel graph. Each graph node is a supervoxel, an atomic group of voxels that is never split (Fig. 2a,b). At any moment in time, the current segmentation is represented by the connected components of the supervoxel graph (Fig. 2c). Two segments can be merged into one by adding an edge to the graph (Fig. 2d), and one segment can be split into two by removing edges (Fig. 2e,f). In order to make splits easy, users can place points on both sides of a proposed split (Fig. 2g) and our proofreading system identifies the edges that need to be removed to separate them. Our system deploys a max-flow min-cut algorithm operating on a local cutout of the supervoxel graph using predicted edge weights as capacities (Fig. 2h).

**Fig. 2.**
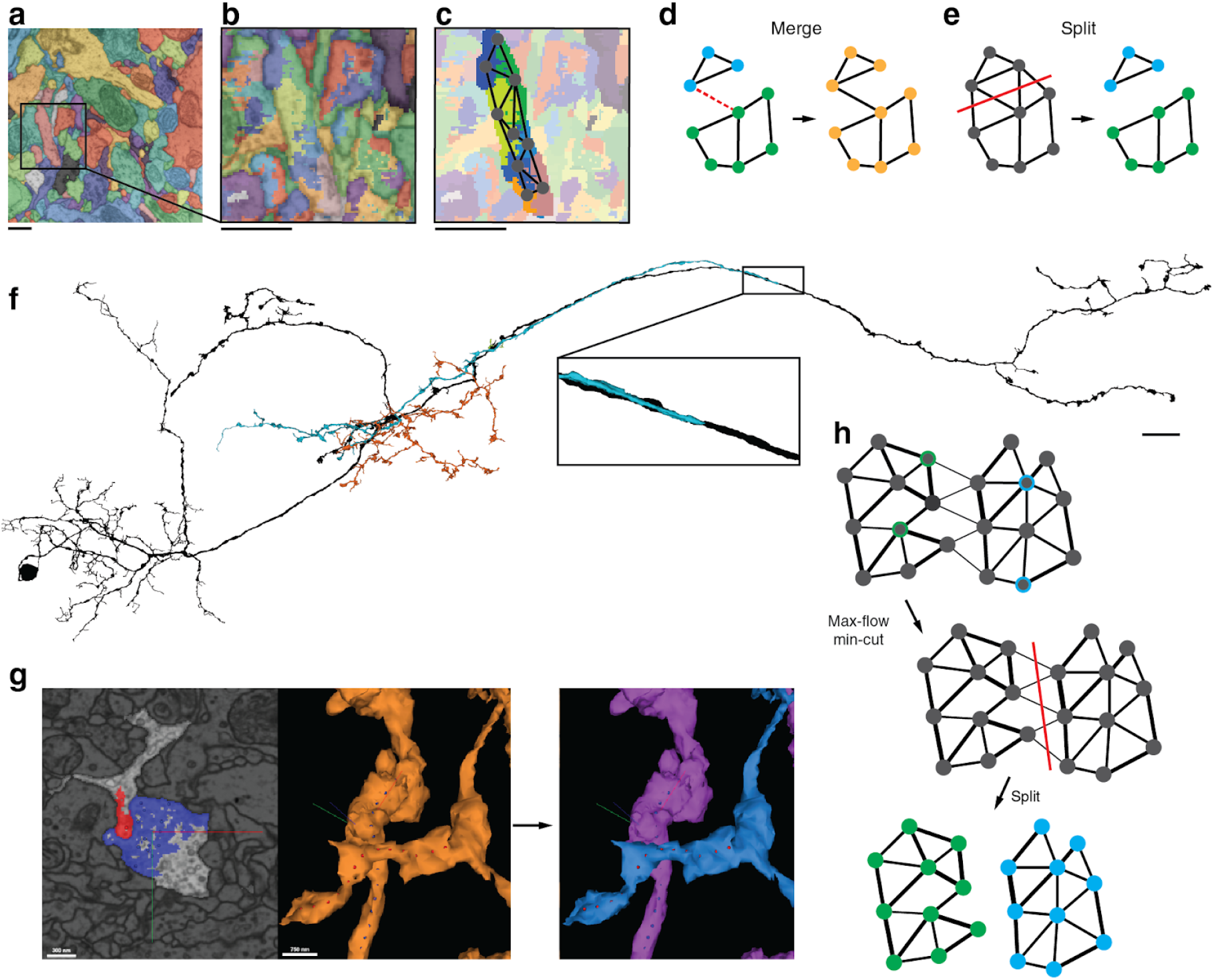
Proofreading the supervoxel graph. (**a**) Automated segmentation overlaid on the EM data. Each color represents an individual putative neuron. (**b**) Each component in (a) is made up of many supervoxels shown in different colors. (**c**) This panel corresponds to the framed square in (a) and the full panel in (b) where we highlighted the supervoxels belonging to a particular neuron and overlaid a cartoon of the supervoxel graph. (**d**) Touching supervoxels (circles) may be connected through edges in the graph indicating that they belong to the same connected component (solid lines). Merge operations add edges between supervoxels resulting in new neuronal components (orange). (**e**) Split operations remove edges resulting in new neuronal components (green, blue). (**f**) Example neuron after proofreading (black). Green, blue and red components were removed during proofreading (see Ext. Data Fig. 4 for the added pieces). While edit operations have global effects, the edits to the supervoxel graph themselves are performed at a local level. (**g**) For splits, users place points (red and blue dots) either in 2D or 3D (center panel) that are linked to the underlying supervoxels (left panel). The proofreading backend then automatically determines which edges need to be removed and performs the split (right panel). (**h**) For the operation shown in (g) the backend performs max-flow min-cut on the local supervoxel graph to determine the optimal cut that separates the user-defined input locations (green and blue framed circles). The thickness of the edges symbolizes the edge weight (cartoon). Scale bars (a,b,c): 1μm (f): 10 μm

Scaling proofreading to a community demands that all users can access the latest state of the segmentation and that multiple users can work on the same neuron without introducing inconsistencies. To allow interactive and continuous proofreading, edits must be resolved quickly and visuals must be updated for the user. At the same time, older states of the segmentation must be accessible for review and publications. However, reads, writes and computations on the supervoxel graph can be time-consuming, because they scale at least linearly with the size of the components. That is because edits have global effects on the connected components even though they only introduce local changes (Fig. 2f). Because of these challenges, no system for community based proofreading of entire neurons exists that scales to datasets as large as FAFB. Existing systems on smaller datasets restrict what proofreaders can work on(J. S. Kim et al. 2014) or do not allow open proofreading by a community (Xu et al. 2020).

We designed a novel data structure, the ChunkedGraph, to address these challenges. The ChunkedGraph leverages the fact that edits only change a small region of a neuron, leaving the rest unchanged. It caches information about connected components spatially allowing it to update components rapidly after edits and restricts the part of the graph that needs to be read and analyzed for them. For this, the nodes of the supervoxel graph are divided into spatial chunks (Extended Data Fig. 3). A supervoxel spanning chunk borders is carved into multiple supervoxels, each contained within a chunk. Each chunk also stores edges between the supervoxels in that chunk. We build an octree on top for storing the connected component information (Fig. 3b). In this tree, nodes in higher layers, abstract nodes, represent connected components in the spatially underlying graph (Fig. 3b-d). Because the ChunkedGraph decouples regions of the same neuron from each other, regions unaffected by an edit do not need to be read and included into calculations, and changes only need to propagate up the tree hierarchy (Fig. 3c, Extended Data Fig. 4). Each segment is a tree, and the ChunkedGraph is a forest of all the segments.

**Fig. 3.**
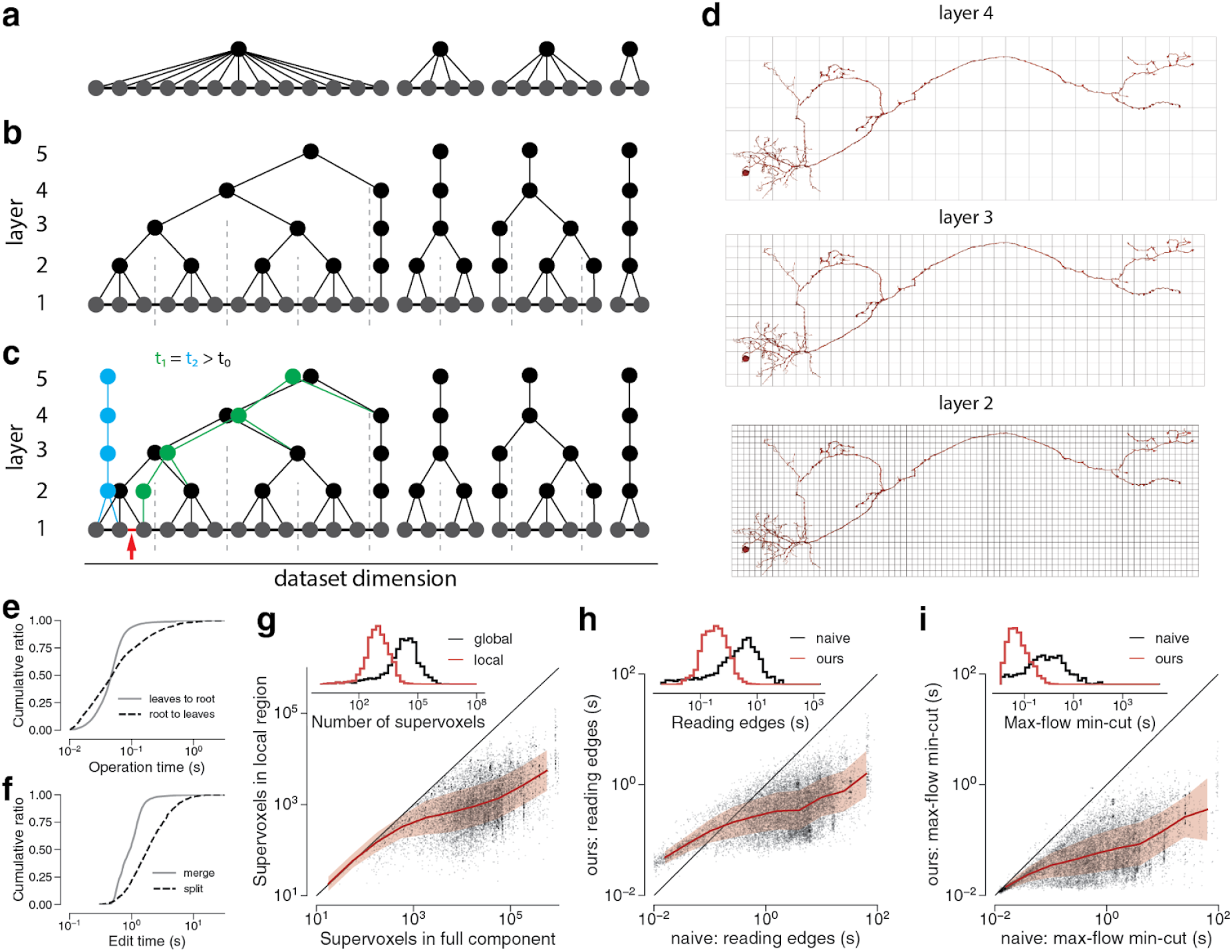
The ChunkedGraph approach for proofreading supervoxel graphs. (**a**) One-dimensional representation of the supervoxels graph. In the simplest approach (naive), one would store the connected component information (neuronal component) in a dedicated parent node. (**b**) We propose an alternative data structure to enable proofreading of very large supervoxel graphs. We store the connected component information in an octree structure where each abstract node (black and colored nodes in levels > 1) represents the connected component in the spatially underlying graph (dashed lines represent chunk boundaries, see Ext. Fig. 3). Nodes on the highest layer represent entire neuronal components. (**c**) Edits to the ChunkedGraph (here, a split; indicated by the red arrow and removed red edge) do not require the loading of the entire supervoxel graph to recompute the neuronal connected components. We can reuse the connected component information stored in the abstract nodes that did not see a change to their underlying graph. (**d**) Chunks double in size along each dimension when going to the higher layer. (**e**) Server response times for the remapping of the connected components from root to supervoxel (N=3,080,494) and supervoxel to root (N=12,096) (**f**) as well as splits (N=2,497) and merges (N=4,612) for real user interactions in the beta-phase of FlyWire. (**g**) Compared to the naive approach (Fig. 3a), our approach reduces the number of supervoxels that need to be loaded for a split (global vs local) (**h**) The subgraph can be read faster and (i) speeds up max-flow min-cut calculations (Fig. 2g,h). The red lines in (g, h, i) are mean and the shaded area standard deviation of bins along x-axis (10 bins); N=15,233.

The ChunkedGraph is initialized by ingesting the initial supervoxel graph created by our automated segmentation pipeline (Dorkenwald et al. 2019). Our pipeline creates supervoxels by grouping voxels that belong to the same cell with very high confidence, according to the affinity predicting neural network. Edges are added to the ChunkedGraph for every pair of neighboring supervoxels in the same segment. Edge weights are also available from the automated segmentation pipeline, and are also ingested into the ChunkedGraph. Proofreading starts from this initial condition, and proceeds by adding and subtracting edges from the ChunkedGraph.

### Visualization of segments in 2D and 3D

FlyWire provides several visualizations to enable the user to find and correct segmentation errors. 2D cross sections of the EM image are displayed in grayscale. Three orthogonal cross sections are available. 2D cross sections of the segmentation are displayed in color, and can be overlaid on the EM images (Extended Data Fig. 5a, left). FlyWire also displays a 3D rendering, a mesh, of user-selected segments (Extended Data Fig. 5a, middle). All of these visualizations utilize Google’s Neuroglancer software (Maitin-Shepard 2020), which enables viewing of volumetric images in a web browser.

When a user interactively selects a supervoxel with a mouse click, the system rapidly displays all supervoxels belonging to the same segment within the field of view of the user. This is accomplished by searching the ChunkedGraph as follows. The search first traverses the tree from the selected supervoxel to the root node at the top level of the hierarchy. For mapping supervoxel to root, the server responded with a median time of 47 ms and 95th percentile of 111 ms (Fig. 3e, n=12,096). Once the search has reached the root, it proceeds back down the tree to identify all supervoxels connected to it within the displayed area, making use of the octree structure of the ChunkedGraph. For mapping root to supervoxels, the server responded with a median time of 48 ms and 95th percentile of 465 ms per displayed chunk (n=3,080,494). Such fast response times are crucial for a globally distributed system if every user is to see the latest state of the segmentation and no data is stored locally. The above times are server response times and were measured during FlyWire’s beta phase (graph with 2.38 billion supervoxels).

### Proofreading by editing the supervoxel graph

Interactive proofreading (Fig. 2g) is implemented using the ChunkedGraph as follows. The user specifies a merge operation by selecting two supervoxels with mouse clicks. An edge between this pair is added to the supervoxel graph (Fig. 2d, Extended Data Fig. 5). Merge edits took 940 ms at median, and 1841 ms at 95th percentile (n=4,612) (Fig. 3f). The user specifies a split operation by selecting supervoxels with mouse clicks (Fig. 2e,g). The system applies a min-cut algorithm to remove a set of edges with minimum weight that leaves the two supervoxels in separate segments (Fig. 2g,h). Split edits had a median time of 1,818 ms, and 95th percentile time of 7,137 ms (n=2,497) (Fig. 3f).

After each edit, the ChunkedGraph generates new abstract nodes in higher layers (> 1, colored nodes in Fig. 2c and Extended Data Fig. 4a). Here, the tree is only traversed in its height and not its width because connected components in neighboring regions are cached in abstract nodes. We use the same abstraction for fast mesh generation of new components by restricting the application of costly and slow meshing algorithms (eg. marching cubes) to single chunks. We only compute meshes from the segmentation for abstract nodes on level 2 (Extended Fig. 3d) and then stitch these to larger components according to the hierarchy such that each abstract node up to a predefined layer has a corresponding mesh. The ChunkedGraph dynamically generates instructions for which mesh files to load for a given component.

We compared the performance of the ChunkedGraph versus an equivalent naive implementation of the supervoxel graph (Methods) (Fig. 3h,i). We measured two different parts of split operation: reading of the edges to compute the split and the min-cut algorithm. The ChunkedGraph greatly benefits from being able to restrict the operations to a subregion (Fig. 3g), leading to orders of magnitude faster reading and calculations (Fig. 3h,i). The ChunkedGraph incurs a minor overhead that is only notable for very small components.

The ChunkedGraph allows concurrent and unrestricted proofreading by many users through serializing edits on a per-neuron level. Edits generate new, timestamped nodes on higher levels (Fig. 3c, Extended Data Fig. 4b), allowing the retrieval of any older state of the segmentation by applying a time filter during tree traversal. Edits can only be applied to the latest version of the segmentation. We implemented the ChunkedGraph with Google’s BigTable (Chang et al. 2008), a low-latency NoSQL database, making use of its timestamping features and atomic row write and read capabilities. For the user, the ability to view the state of a cell from any past time point in the proofreading process is helpful for reviewing one’s own work or the work of others. This is analogous to viewing past versions of a Wikipedia article, which are recreated using the edit history (Priedhorsky et al. 2007) (Fig. 4a).

**Figure 4.**
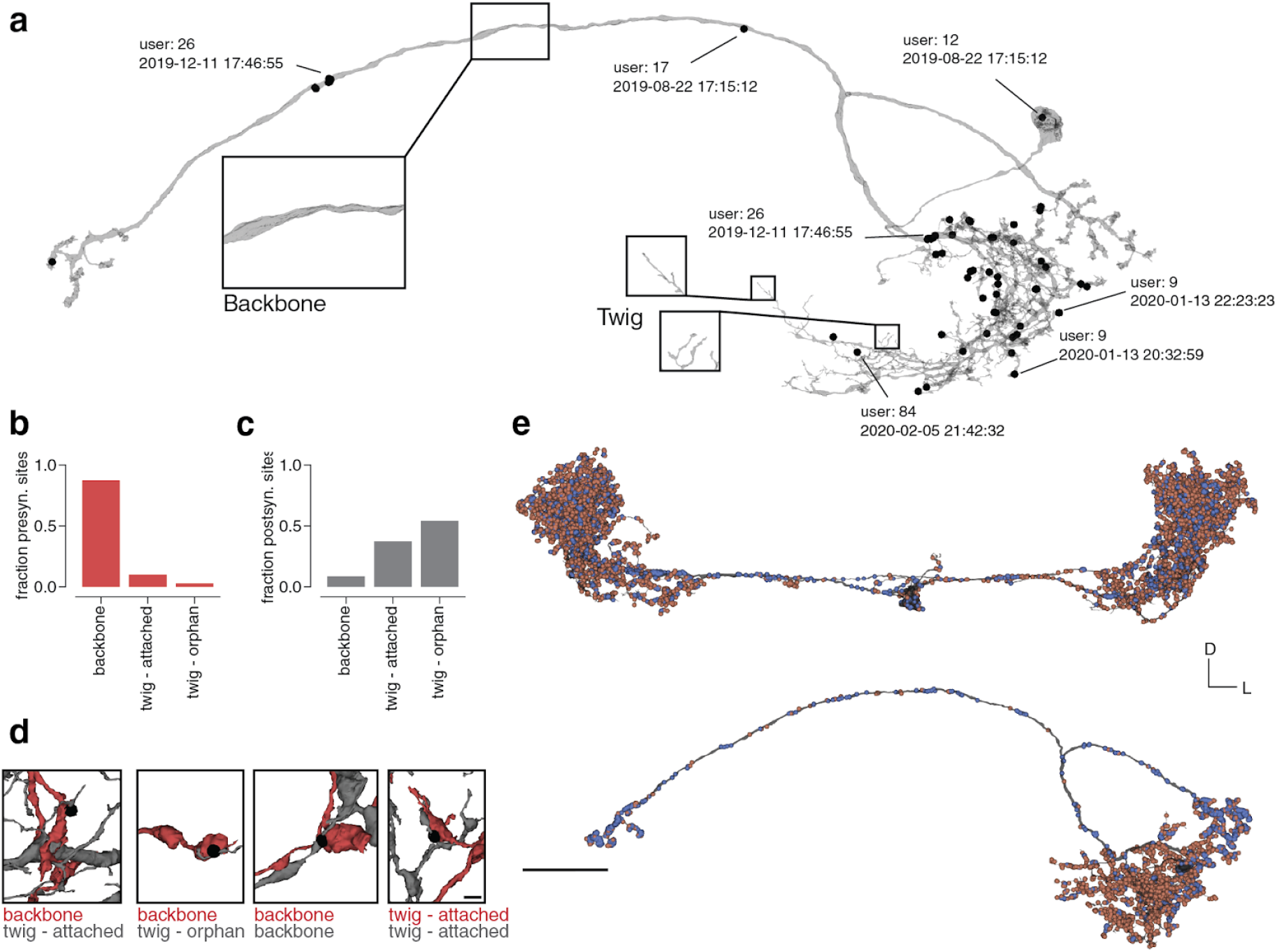
Attaching automatically detected synapses to neurons. (**a**) Each edit (black dot) is linked to a user and timestamp enabling the retrieval of the edit history and credit assignment post-hoc. Neurons in the fly brain can be roughly partitioned into sections known as “twigs” and “backbone”. (**b**) We classified pre- and (**c**) post-synaptic segments based on their morphology and whether they are attached to a bigger component that will be attached during a conservative procedure. (**d**) shows examples of these assessments. (**e**) shows a proofread WV-WV PN (top) and AMMC-B1 (bottom) with automatically detected synapses attached (blue: presynaptic, red: postsynaptic). Scale bar: (d): 1 μm; (e) 25 μm

### Extracting synaptic connections

With hundreds of millions of synaptic connections in the fly brain (Buhmann et al. 2019), automated synaptic partner identification is required to facilitate connectivity analysis at scale. Several methods have been proposed for synapse detection in large EM datasets (Heinrich et al. 2018; Staffler et al. 2017; Dorkenwald et al. 2017; Huang, Scheffer, and Plaza 2018; Buhmann et al. 2019; Turner et al. 2019) but only a few solved the unique problem of partner assignments in polyadic synapses in the fly (Buhmann et al. 2018; Buhmann et al. 2019; Huang, Scheffer, and Plaza 2018; Kreshuk et al. 2015). FlyWire is compatible with any existing and future method that identifies synaptic partners and their pre- and postsynaptic sites. Currently, only Buhmann et al.(Buhmann et al. 2019) demonstrated their method on the whole fly brain. We imported their synapses into our realigned coordinate space and made them available to the community.

A fly neuron is said to consist of a thicker, microtubule-rich “backbone” and numerous thin “twigs” (Schneider-Mizell et al. 2016) (Fig. 4a). The distinction can be somewhat subjective in borderline cases, but is quite useful in practice. The automated segmentation contains many small “orphan twigs” that are not assigned to any large neuronal object. Attaching orphan twigs is time consuming and difficult because twigs contain very thin processes. Therefore, our proofreaders did not correct orphan twigs for the most part. The hemibrain similarly contains many orphan twigs after proofreading (Xu et al. 2020).

Avoiding the excessive labor of reattaching twigs comes at some cost: synapses involving orphan twigs will be missing from the reconstruction. Fortunately, many fly neurons are redundantly connected, with up to hundreds of synapses possible between a connected pair (Takemura et al. 2013). It has been proposed that a neuron pair sharing fewer than 10 synapses represents a minor connection (Meinertzhagen 2018). If omissions of synapses are statistically independent, then connections will be recalled with a probability that increases with the number of synapses involved (Schneider-Mizell et al. 2016).

We quantified the number of missing synapses due to orphan twigs by evaluating the segmentation at 612 randomly picked synaptic locations from Buhmann et al. (see Methods). For each of these synapses an expert judged whether the pre- and postsynaptic reconstructions were at a backbone or twig and whether the twig was attached (“twig - attached”) to a backbone or orphan (“twig - orphan”) (Fig. 4b-d, see Methods). We found that 40.6% of all postsynaptic and 78.2% of presynaptic twigs were attached to backbones. In aggregate, we expect our proposed conservative proofreading to at least include all “backbone” and “twig - attached” segments in a proofread neuron. Our assessment leads to a combined estimate of 44.6% of synapses with pre- and postsynaptic segments attached on average after proofreading. In practice, major connections (>9 synapses, 99.7% with at least one synapse) and most minor connections with at least 3 synapses are maintained (83% with at least one synapse) due to the redundant nature of connections in the fly brain (Schneider-Mizell et al. 2016; Meinertzhagen 2018).

For analysis, we assign synapses to neurons based on their pre- and postsynaptic coordinates (Fig. 4e). Since proofreading changes the assignment of synapses to neurons, we continuously release new versions of the synapse table.

### Quantification of proofreading effort and accuracy

To assess effort required to proofread neuronal backbones, we proofread 183 neurons mostly with projections in early mechanosensory neuropils (antennal mechanosensory and motor center (AMMC), wedge (WED), and ventrolateral protocerebrum (VLP)). Each neuron was proofread by three different people in successive rounds. The number of errors corrected decreased after the first round (Fig. 5a), most notably large corrections (volumetric difference > 1μm^3^) decreased from a median of 8 in the first round to medians of 1 and 0 in the second and third round (Fig. 5b).

**Figure 5.**
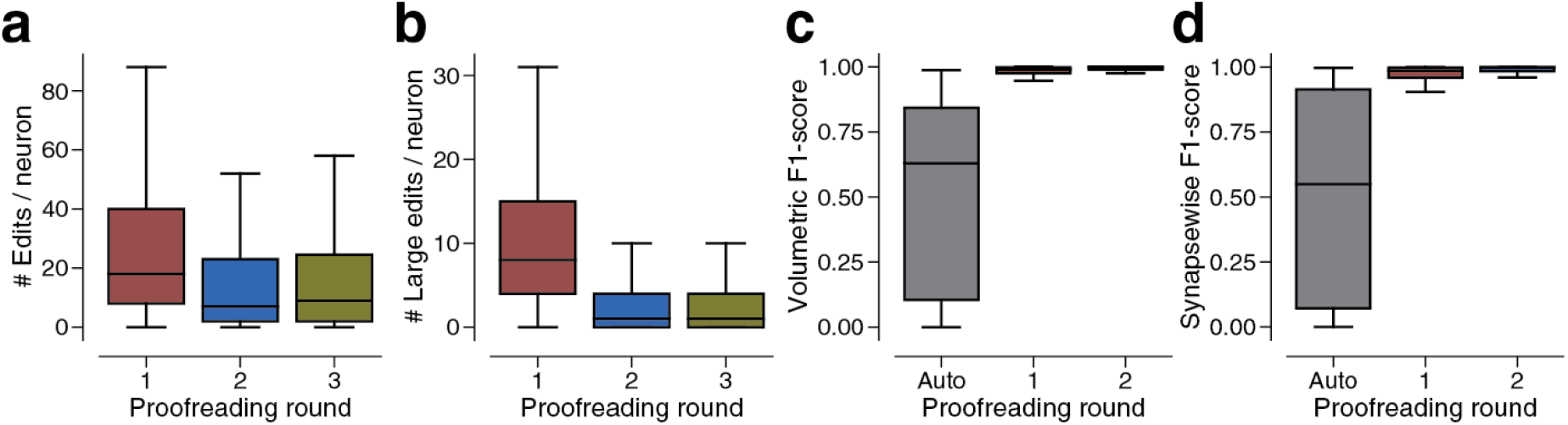
Proofreading in FlyWire. (**a**) Number of edits per neuron (N=183) and proofreading round. (**b**) Same as (a) but restricted to large edits (> 1μm^3^). Large edits are usually edits on “backbones” whereas small edits tend to affect “twigs”. (**c**) Comparing the F1-Scores (0-1, higher is better; with respect to proofreading results after three rounds) between different proofreading rounds according to volumetric completeness and (**d**) assigned synapses. “Auto” refers to reconstructions without proofreading.

To quantify the impact of the different proofreading rounds further, we scored the reconstructions after each round with respect to their state after the third round. We calculated F1-Scores with respect to volumetric completeness and correct synapse assignments (pre- and postsynaptic irrespectively) (Fig. 5c,d). One round of proofreading was enough to recover an accurate morphology and synapse assignment in most cells (median F1-Scores of .99 for both volumetric and synapse based assessments). We measured times for 49 of the 183 neurons and found that first-round proofreading took a median of 90 minutes per neuron (mean: 135 minutes).

Researchers can use FlyWire to proofread to their desired level of accuracy and some researchers have reported scientific benefits without any proofreading at all. For others it may be sufficient to proofread backbones, as in Deutsch et al. (Deutsch, Pacheco, and Encarnacion-Rivera 2020) using FlyWire to reconstruct a circuit involved in female social behavior.

### Novel connections and subtypes in mechanosensory pathways

To validate FlyWire as a circuit discovery platform, we proofread and analyzed 180 neurons (belonging to seven major types) in the AMMC, WED, and VLP neuropils (Fig. 6a), all previously shown to respond to mechanosensory stimuli, such as sound or wind (Tootoonian et al. 2012; Lai et al. 2012; Vaughan et al. 2014; Yamada et al. 2018). These neurons were found based on their previously identified morphology and cell body location (Kamikouchi et al. 2009; Lai et al. 2012; Tootoonian et al. 2012) (see Methods, and Supplementary Table 1 for numbers of neurons per cell type).

**Figure 6.**
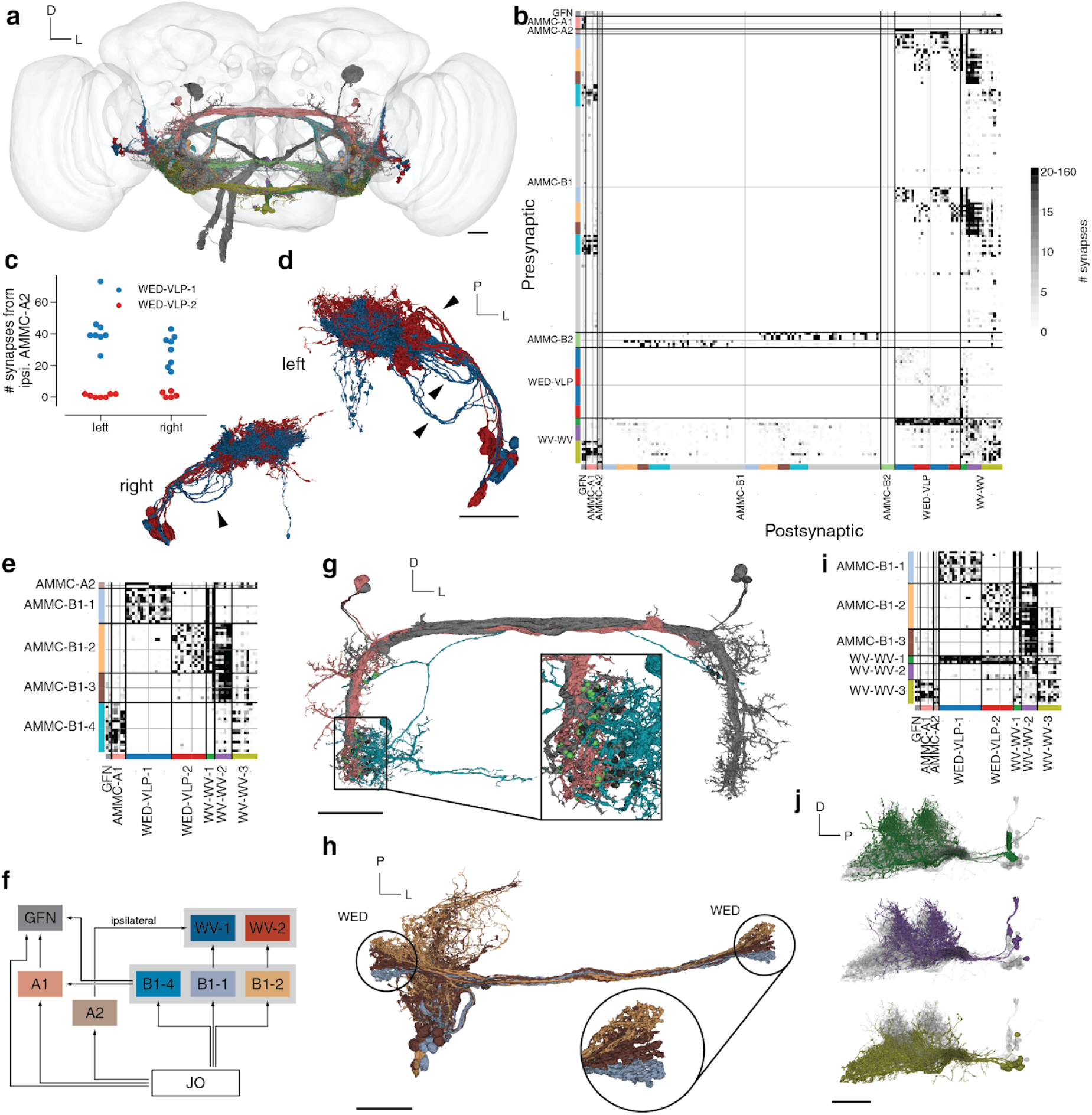
Auditory connectivity extracted with FlyWire. (**a**) We included 180 neurons from both hemispheres into our analysis, here colored by their cell type (black: giant fiber neuron, turquoise: AMMC-A1, orange: AMMC-A2, dark blue: AMMC-B1, light blue: AMMC-B2, red: WED-VLP PN, green: WV-WV PN). (**b**) Connectivity diagram between all 180 neurons ordered by cell type. Gray through lines divide cells from different hemispheres (left/top: left hemisphere, right/bottom: right hemisphere) and colored bars separate putative sub-celltype (see Extended Data Fig. 6 for matrices with different colormap cutoffs). (**c**) Analyzing the synaptic inputs of the WED-VLP neurons onto the suggested groups shows differential input from AMMC-A2 subtypes. (**d**) The two WED-VLP groups differ in their morphology (see arrow heads). (**e**) We grouped AMMC-B1 neurons according to their outputs on to other cell types in the AMMC. (**f**) Abridged summarized connectivity diagram showing the selective targeting of WED-VLP PNs (WV) by AMMC-B1s (B1) and the cross-pathway innervation by AMMC-B1-4s. (**g**) The connectivity based groupings of AMMC-B1 neurons correspond to a stration of axonal arbors in both hemispheres (arrow heads). (**h**) One group of AMMC-B1 neurons targets AMMC-A1 neurons in both hemispheres (red: AMMC-A1, turquoise: AMMC-B1-4, gray: all AMMC-A1s). We automatically detected 217 synapses from this specific AMMC-B1 neuron onto all AMMC-A1 neurons (balls), 87 of which on the shown AMMC-A1 (green balls). (**i**) Commissural neurons vary in their connectivity with other cell types. (**j**) The suggested subgrouping correlates with collocations of their cell bodies as well as different arborizations. Scale bars: 25 μm

Airborne mechanosensory stimuli activate Johnston’s Organ (JO) of which distinct populations send broadly tonotopic projections to five different zones within the AMMC. AMMC neurons in turn connect with the WED and VLP (Kamikouchi et al. 2009). We identified neurons proposed to have dendrites in AMMC zones A (AMMC-A1, AMMC-A2, GFN) and B (AMMC-B1, AMMC-B2) and therefore receive inputs largely from JO-As and JO-Bs respectively (H. Kim et al. 2020). Although prior work identified only 10 AMMC-B1 neurons per hemisphere (H. Kim et al. 2020; Lai et al. 2012), we identified 61 neurons in the left hemisphere and 58 neurons in the right hemisphere, all with clear B1 morphology (Extended Data Fig. 6). We additionally identified neurons belonging to cell types WED-VLP (aka iVLP-VLP (Lai et al. 2012)) and WV-WV (aka iVLP-iVLP (Lai et al. 2012) or WED-WED (Clemens et al. 2015)).

AMMC-B1 neurons respond strongly to frequencies present in conspecific courtship songs (Azevedo and Wilson 2017) and are thought to target WED-VLPs, based on the proximity of their processes (Lai et al. 2012), forming a putative pathway for courtship song processing in the brain. GFNs and AMMC-A1 neurons on the other hand, while responsive to song stimuli (Tootoonian et al. 2012; Clemens et al. 2015), are core components of the *Drosophila* escape pathway (von Reyn et al. 2014; Allen et al. 2006). We determined if there was any overlap between these two pathways and also looked for important subtypes, based on connectivity and morphology, within each cell type. Finally, as FlyWire enables identification of synaptic connections across both brain hemispheres, we found pathways that could mediate the detection of sound or wind direction via cross-hemispheric connections. To do this, we created a wiring diagram between all 180 identified neurons by combining our reconstructions (Fig. 6a) with the synapses from Buhmann et al. (Buhmann et al. 2019) with minor additional synapse proofreading (Fig. 6b, see Ext. Data Fig. 7 for a matrix ordered by subtype and a matrix with different colormap threshold, see Methods).

Our analysis confirms previously proposed pathways between AMMC-A1 and GFN (Phelan et al. 2008) as well as AMMC-B1 and WED-VLPs (Lai et al. 2012). However, we found that only a minority of the AMMC-B1 neurons innervated WED-VLPs (left: 15/61, right: 14/58, Supplementary Table 1, Fig. 6b) and that two subgroups of AMMC-B1s targeted two subgroups of WED-VLPs. This partition of WED-VLPs was directly related to input from ipsilateral AMMC-A2s (Fig. 6c) and a morphological separation of their arbors (Fig. 6d). It stands out that WED-VLP-1 neurons receive convergent input from AMMC-A2 and AMMC-B1-1 neurons, positioning them to encode both auditory stimulus motion energy and sound vibration (Azevedo and Wilson 2017).

AMMC-B1 neurons can be divided into at least 5 subtypes (see Extended Data Fig. 6 for a morphological comparison and Supplementary Table 2 links to the data) - AMMC-B1-1 and AMMC-B1-2 neurons project to WED-VLP neurons, AMMC-B1-3 neurons target only the WV-WV neurons, and AMMC-B1-4 neurons send outputs (with up to 90 synapses per connection) to the GFN and AMMC-A1 neurons, suggesting the existence of (previously unknown) feedback from the courtship song processing pathway to the escape pathway (Fig. 6h). AMMC-B1-u (u for unidentified) neurons had almost all of their synapses onto neurons not included in our set of 180 neurons. When validating these subtypes morphologically, we found that the axonal arbors of AMMC-B1-1, -2 and -3 striate the WED in both hemispheres (Fig. 6g). AMMC-B2 neurons receive input from ipsilateral JO-B neurons, are GABAergic, and proposed to sharpen the tuning of AMMC-B1 for sound frequencies (Yamada et al. 2018); we found that they only target AMMC-B1 neurons in the contralateral hemisphere (Fig. 6b, Extended Fig. 1a), suggesting a role in spatial localization of sounds.

WV-WVs are GABAergic (Lai et al. 2012) with cell bodies in the center of the brain and symmetrical processes in both hemispheres - these neurons are therefore well positioned to provide feedback inhibition within the circuit. We identified a subgroup that targets GFN, AMMC-A1 and AMMC-A2 neurons in both hemispheres (WV-WV-3) as well as a subgroup that strongly synapses onto WED-VLPs (WV-WV-1). Lastly, we identified a group (WV-WV-2) receiving input predominantly from AMMC-B1-2 and AMMC-B1-3 neurons but not from AMMC-B1-1 neurons (Fig. 6i). That WV-WV-3s receive input from AMMC-B1-4 neurons and contact GFN and AMMC-A1 neurons is again consistent with cross-talk between the courtship song and escape pathways. When observing the morphology of these three types of WV-WV neurons, we found a correlation with the location of cell bodies and arborizations (Fig. 6j).

In sum, this analysis highlights the value of mapping connections across both brain hemispheres and supports the utility of EM connectomics in both finding novel links between (previously considered distinct) pathways and identifying important distinctions within cell types known from light microscopy studies.

### Community organization

Users are currently being recruited from *Drosophila* labs. We are starting with professional scientists, who are inherently incentivized for productivity and accuracy, because their own discoveries depend on their proofreading. Later on, we plan to expand recruitment to non-scientist volunteers. This is more complex, because “gamification” will be required to incentivize users (J. S. Kim et al. 2014).

During onboarding, users study self-guided training materials (see “Training Materials” on https://flywire.ai), and practice proofreading in a “Sandbox” dataset. Users are granted proofreading privileges in the real dataset after passing an entry test. In Wikipedia, unqualified or malicious users may introduce mistakes into articles. However, even without tests the completeness and accuracy of articles in Wikipedia tends to increase over time as users detect and correct omissions or errors in articles. Some studies have found that Wikipedia is approximately as accurate as traditional encyclopedias (Giles 2005). FlyWire utilizes the same basic mechanism of crowd wisdom as Wikipedia, iterative collaborative editing, while adding a safety layer through entry-level testing and subsequent spot checks of proofreading quality.

Members must consent to follow the FlyWire community principles (https://flywire.ai). As mentioned earlier, the most important principle is openness. Users may access information about any cell, such as cell type, proofreading status, or who made edits. Another important principle is fairness. Every edit to a neuron is regarded as a “micropublication,” and the identity of the authoring member is recorded in the database along with the edit. Members agree to freely license their edits to others provided their contributions are attributed. When using FlyWire reconstructions in a scientific publication or presentation, users must obtain the agreement of other significant contributors, and make their neurons “public”. Once “public” neurons are available to anyone (such as the neurons included in this publication). Such credit assignment procedures attempt to make FlyWire fair while maintaining its openness.

## DISCUSSION

FlyWire is an implementation of our proposal for an open community to proofread an automated reconstruction of the entire *Drosophila melanogaster* brain. Interactive proofreading in FlyWire is enabled by a novel data structure for proofreading large connectomics datasets, the ChunkedGraph. As a demonstration of the segmentation and proofreading tools in FlyWire, we reconstructed 180 known mechanosensory neurons across both hemispheres and introduced new subdivisions of known cell types based on their connectivity and morphology. Further, we found new connections between the courtship song and escape pathways. Our results were corroborated by their replication in the other hemisphere. FlyWire’s completeness of the brain allows researchers to identify all partners of a neuron. While we observed a striking hemispheric symmetry in the connectivity between the analyzed mechanosensory neurons, we also found asymmetric interactions of neurons crossing the midline, providing a potential mechanism for spatial localization of sounds, a challenging problem for flies with closely-spaced antennal auditory receivers (Morley et al. 2012).

It is likely that each whole brain connectome will require proofreading by many people for years, in spite of increases in the accuracy of automated reconstruction. We propose that whole-brain connectomics for each animal species could benefit from a decentralized approach that crowdsources proofreading to the researchers of that species. This approach would make circuits available with zero delay, accelerating research. Researchers would be able to prioritize proofreading of their own circuits of interest, and researchers could choose to proofread to any accuracy level required by their own scientific questions.

Using the current segmentation’s mean backbone proofreading time of approximately 135 minutes per neuron, and an estimate that the *Drosophila* brain contains approximately 100,000 cells, a whole-brain connectome of these backbones with their existing twigs would require 112.5 person-years of proofreading assuming the use of automatic synapse detection. Ongoing improvements in both the automatic segmentation and the proofreading interface will reduce the number of errors further and make it possible to find and correct the remaining ones more rapidly. Proofreading can be sped up by having the computer automatically identify likely errors and suggest corrections (Zung et al. 2017).

At this writing, over 100 researchers from over 30 labs have been onboarded and trained for FlyWire, and membership is expanding. There are hundreds of labs studying *Drosophila* neural circuits worldwide, and the *Drosophila* research community has a long history of sharing and collaboration. Furthermore, the automated segmentation is now so accurate that interesting science can be discovered by only modest proofreading effort.

---

**Extended Data Figure 1.**
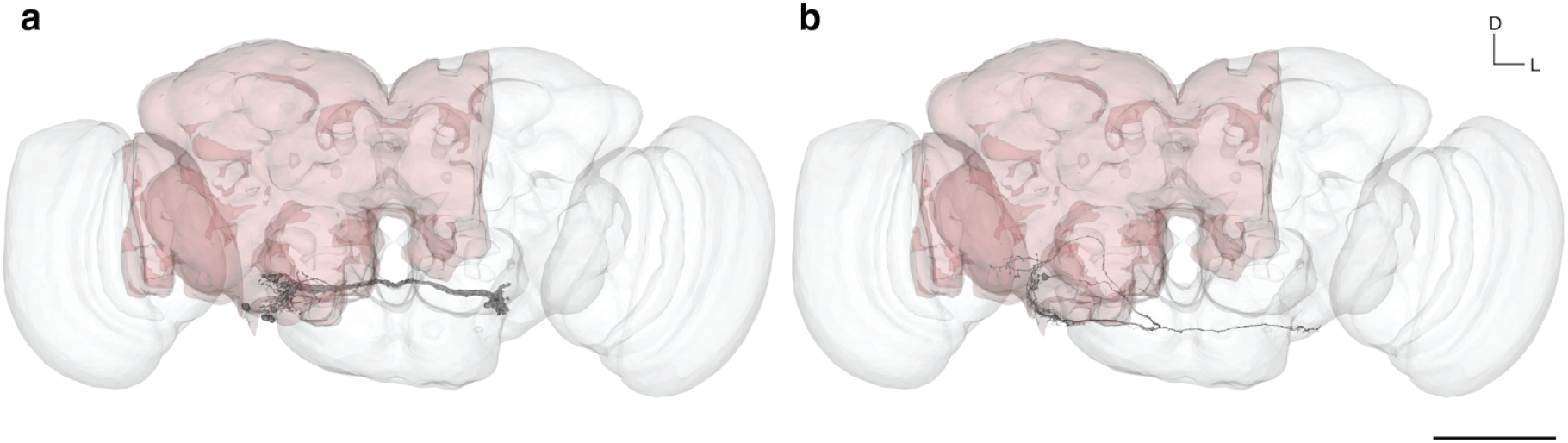
Full brain rendering and comparison with the hemibrain. A neuropil rendering of the fly brain (white) is overlaid with a rendering of the hemibrain (see Methods) and proofread reconstructions of neurons from the antennal mechanosensory and motor center (AMMC). The proofread reconstructions of all (**a**) AMMC-B2 neurons from the right hemisphere and the (**b**) the AMMC-A2 neuron from the right hemisphere are added, illustrating how many neurons are only partially contained in the hemibrain dataset. Scale bar: 100 μm

**Extended Data Figure 2.**
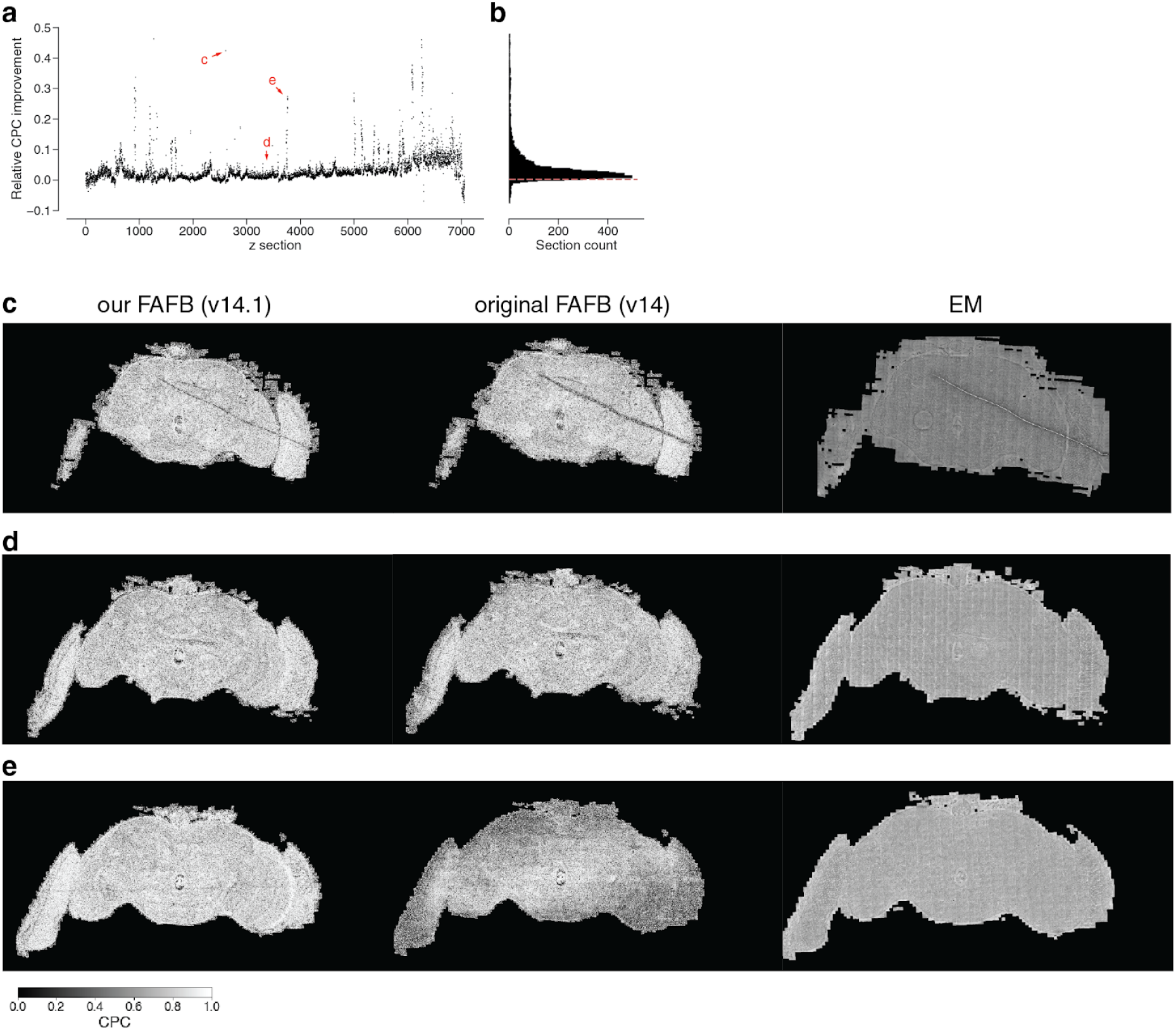
Quality of EM image alignment. We calculated the chunked pearson correlation (CPC) between two neighboring sections in the original alignment (v14) and our re-aligned data (v14.1). (**a**) The relative change of CPC between the original and our re-aligned data per section (**b**) is almost always positive (dashed red line is at 0). (c, d, e) show examples from (a) where (**c**) the CPC improved through a better alignment around an artifact, (**d**) the CPC is almost identical and (**e**) the CPC overall improved due to a stretch of poorly aligned sections in the original data that were resolved in v14.1.

**Extended Data Figure 3.**
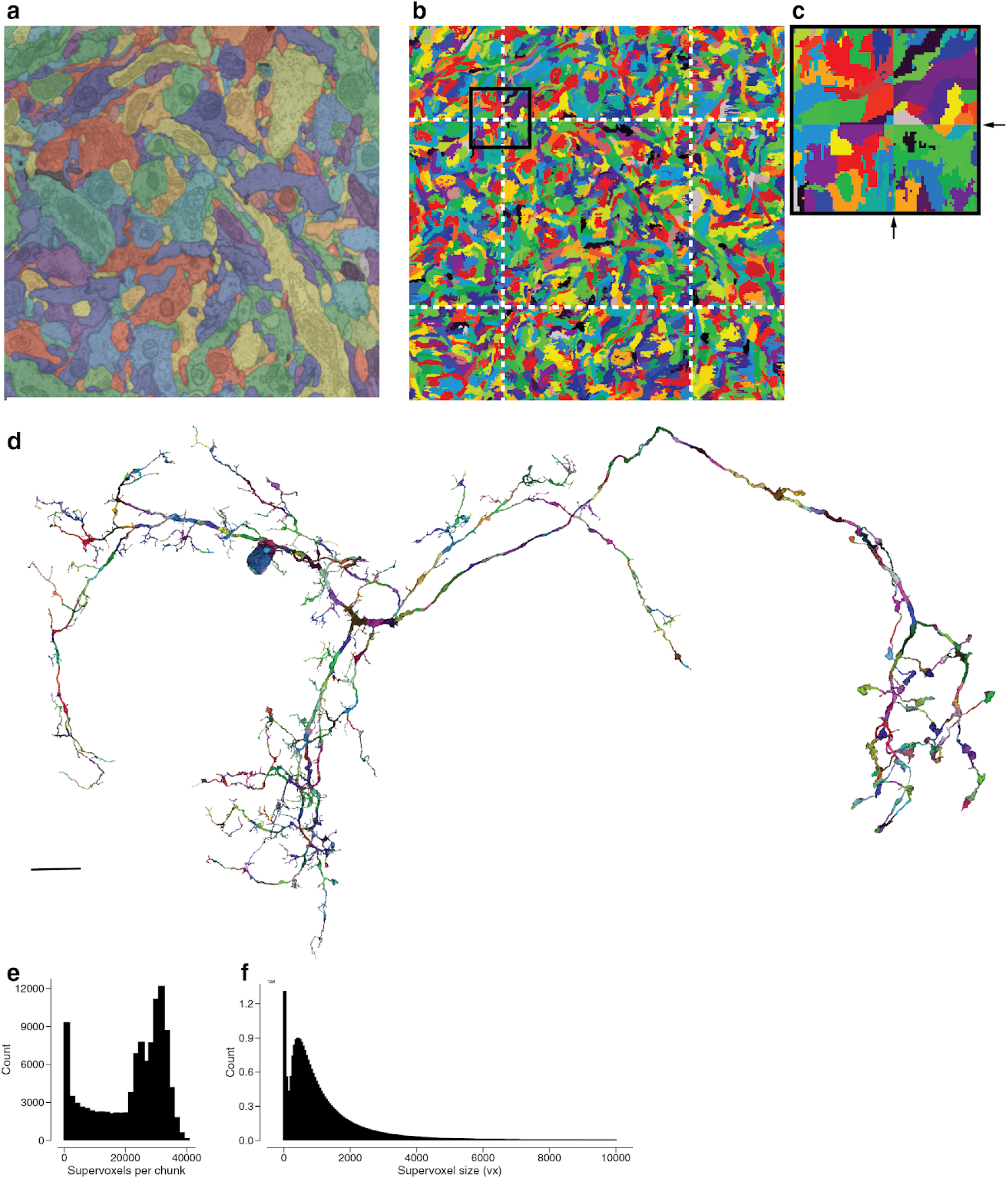
Chunking the dataset. (**a**) Automated segmentation overlayed on the EM data. Each different color represents an individual putative neuron. (**b**) The underlying supervoxel data is chunked (white dotted lines) such that each supervoxel is fully contained in one chunk. (**c**) A close up view of the box in (b). (**d**) We apply the same chunking scheme to the meshes, requiring only minimal mesh recomputations after edits. (**e**) The irregular shape (not a box) of the fly brain leads to a high diversity of the number of supervoxels in each chunk (median: 25661). (**f**) The median supervoxel contains 792 voxels. All very small supervoxels (< 200 voxels) are the result of chunking.

**Extended Data Figure 4.**
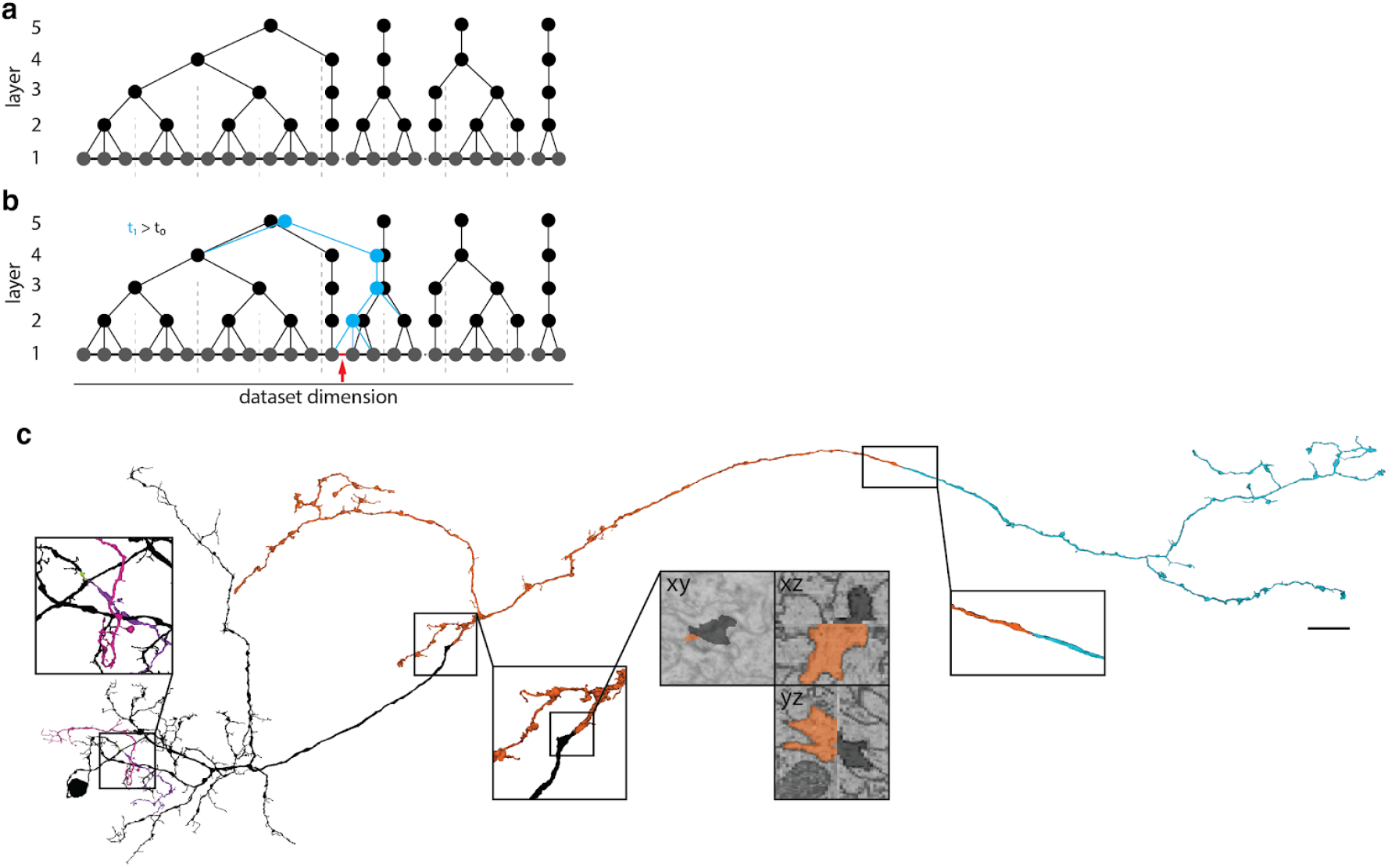
Proofreading with the ChunkedGraph. (**a, b**) Analogous to the split operation in Fig. 3a, b a merge operation does not require the loading and writing to and from the entire connected component since we can reuse the connected component information stored in the abstract nodes that did not see a change to their underlying graph. (**c**) During proofreading, all shown segments were added together. This is the same neuron for which removed segments were shown in Fig 2. Scale bar (c): 10 μm

**Extended Data Figure 5.**
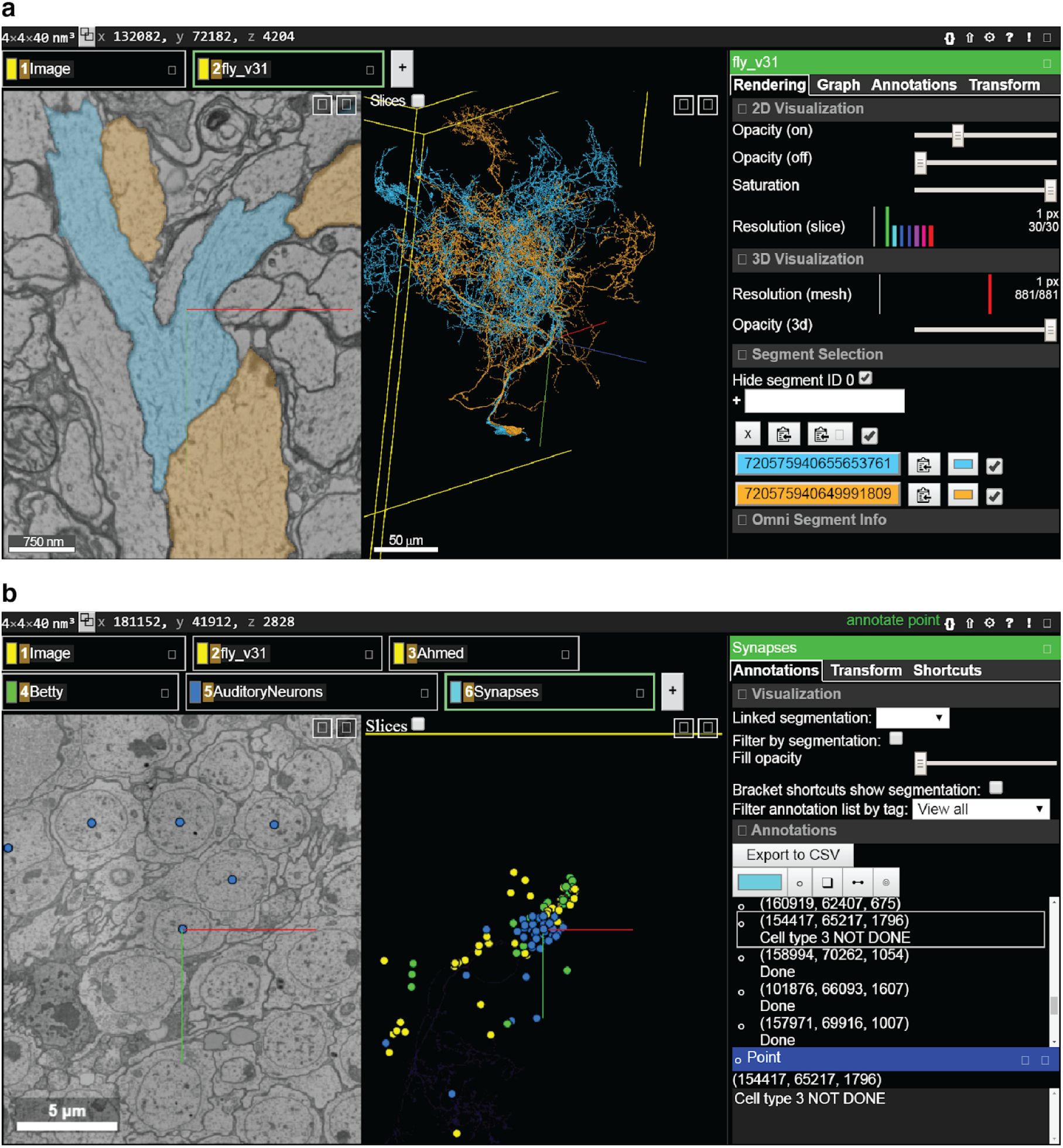
The FlyWire proofreading platform. (**a**) The most common view in FlyWire displays the EM image in grayscale overlaid with segmentation in color (left panel), a 3D view of selected cell segments (middle), and menus with multiple proofreading tools (right). (**b**) Annotation tools include points, which can be used for a variety of purposes such as marking particular cells or synapses.

**Extended Data Figure 6.**
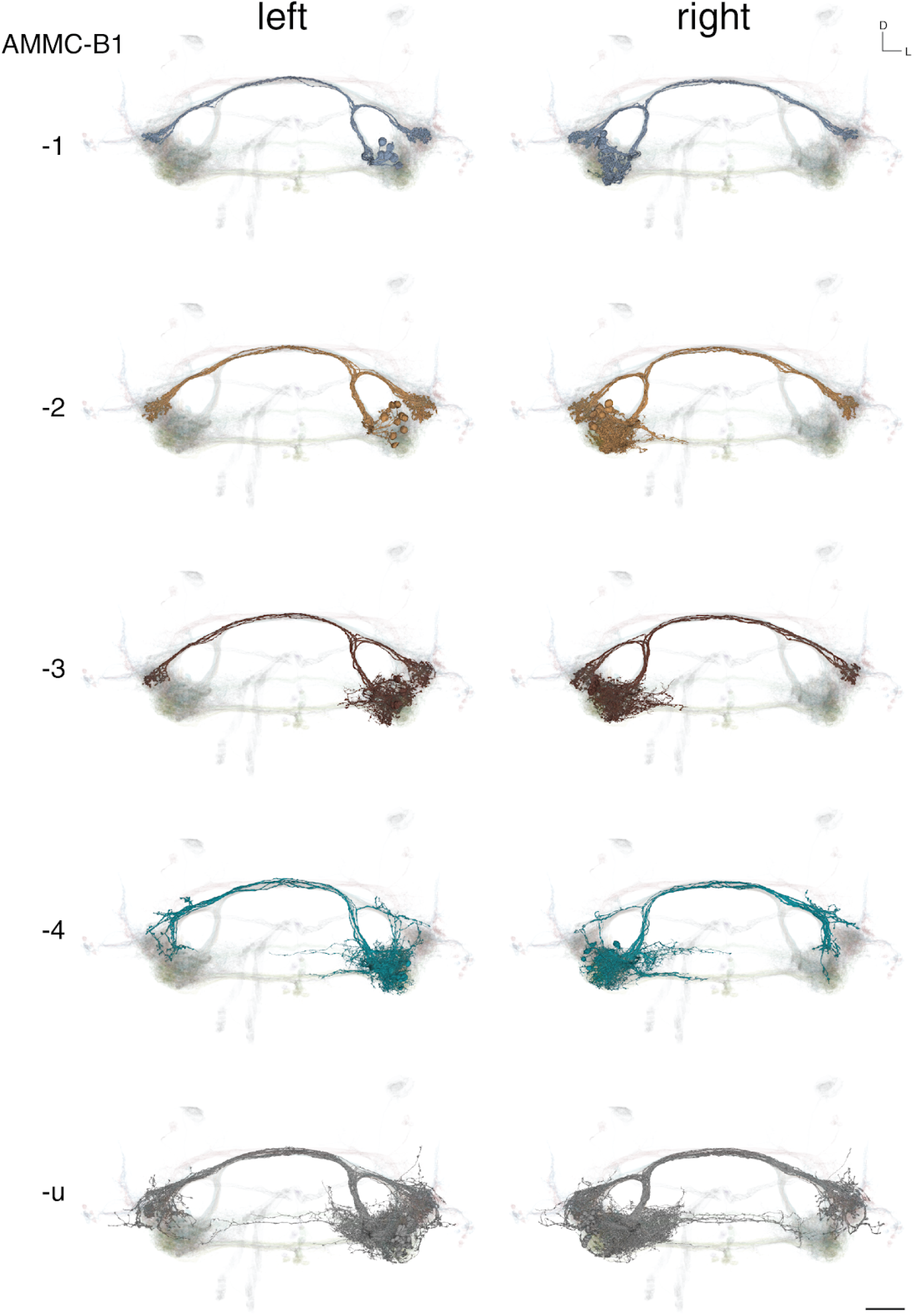
Renderings of AMMC-B1 subtypes. grouped by subtype and hemisphere. All 180 neurons included in the mechanosensory analysis are shown as backdrop. Scale bar: 25 μm

**Extended Data Figure 7.**
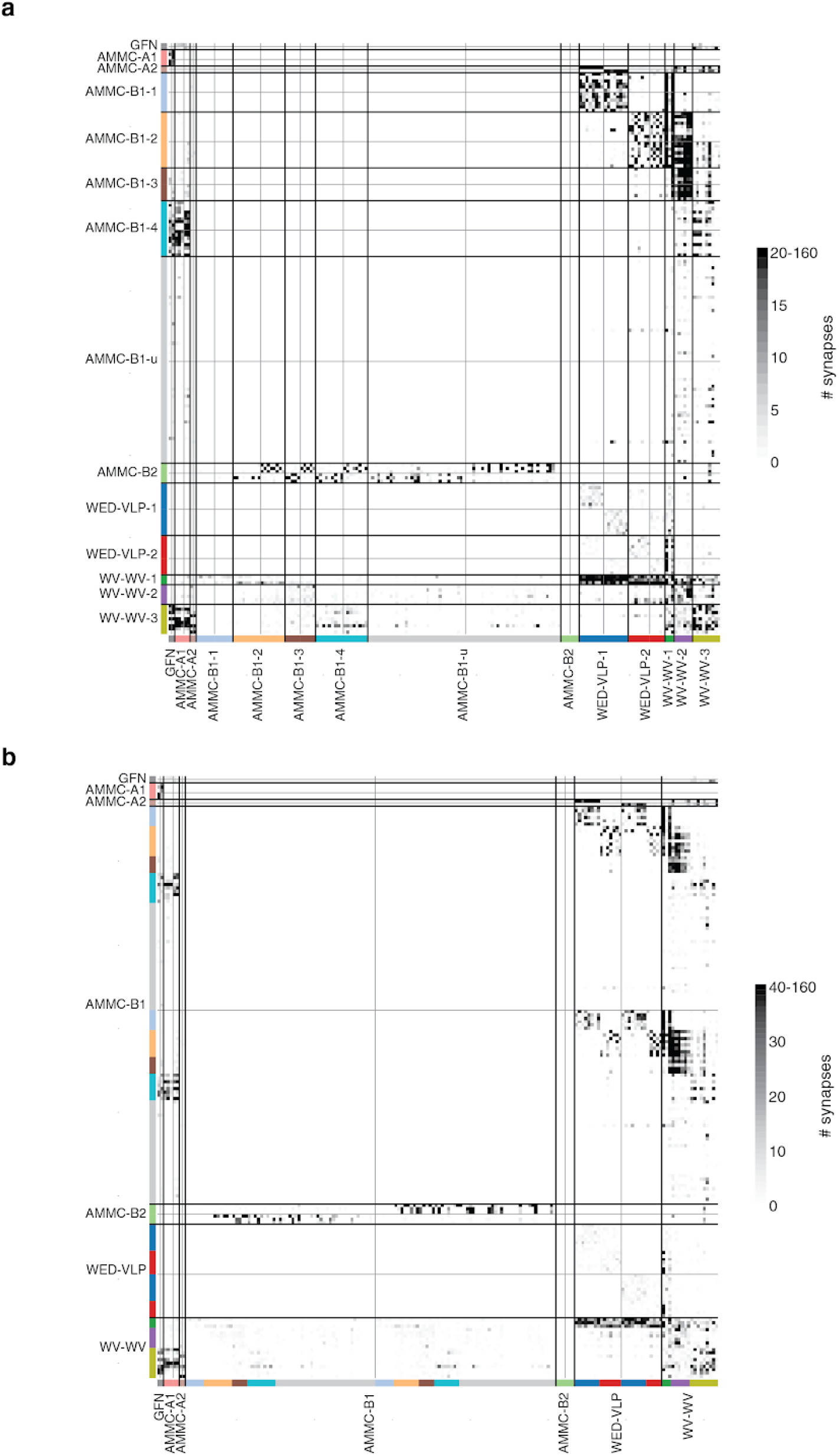
Connectivity diagrams. (**a**) Diagram from Figure 6b reordered by subtype (**b**) Same diagram as in Figure 6b with different colormap threshold.

**Supplementary Table 1.** Cell type counts for mechanosensory analysis.

| Cell type | Subtype | Hemisphere | Count | Total count |
| --- | --- | --- | --- | --- |
| AMMC-A1 | - | left | 3 | 5 |
|  | - | right | 2 |  |
| AMMC-A2 | - | left | 1 | 2 |
|  | - | right | 1 |  |
| AMMC-B1 | 1 | left | 6 | 119 |
|  |  | right | 6 |  |
|  | 2 | left | 9 |  |
|  |  | right | 8 |  |
|  | 3 | left | 5 |  |
|  |  | right | 5 |  |
|  | 4 | left | 9 |  |
|  |  | right | 8 |  |
|  | u | left | 32 |  |
|  |  | right | 31 |  |
| AMMC-B2 | - | left | 3 | 6 |
|  | - | right | 3 |  |
| GFN | - | left | 1 | 2 |
|  | - | right | 1 |  |
| WED-VLP | 1 | left | 8 | 28 |
|  |  | right | 8 |  |
|  | 2 | left | 7 |  |
|  |  | right | 5 |  |
| WV-WV | 1 | unpaired | 3 | 18 |
|  | 2 | unpaired | 6 |  |
|  | 3 | unpaired | 9 |  |

**Supplementary Table 2.**
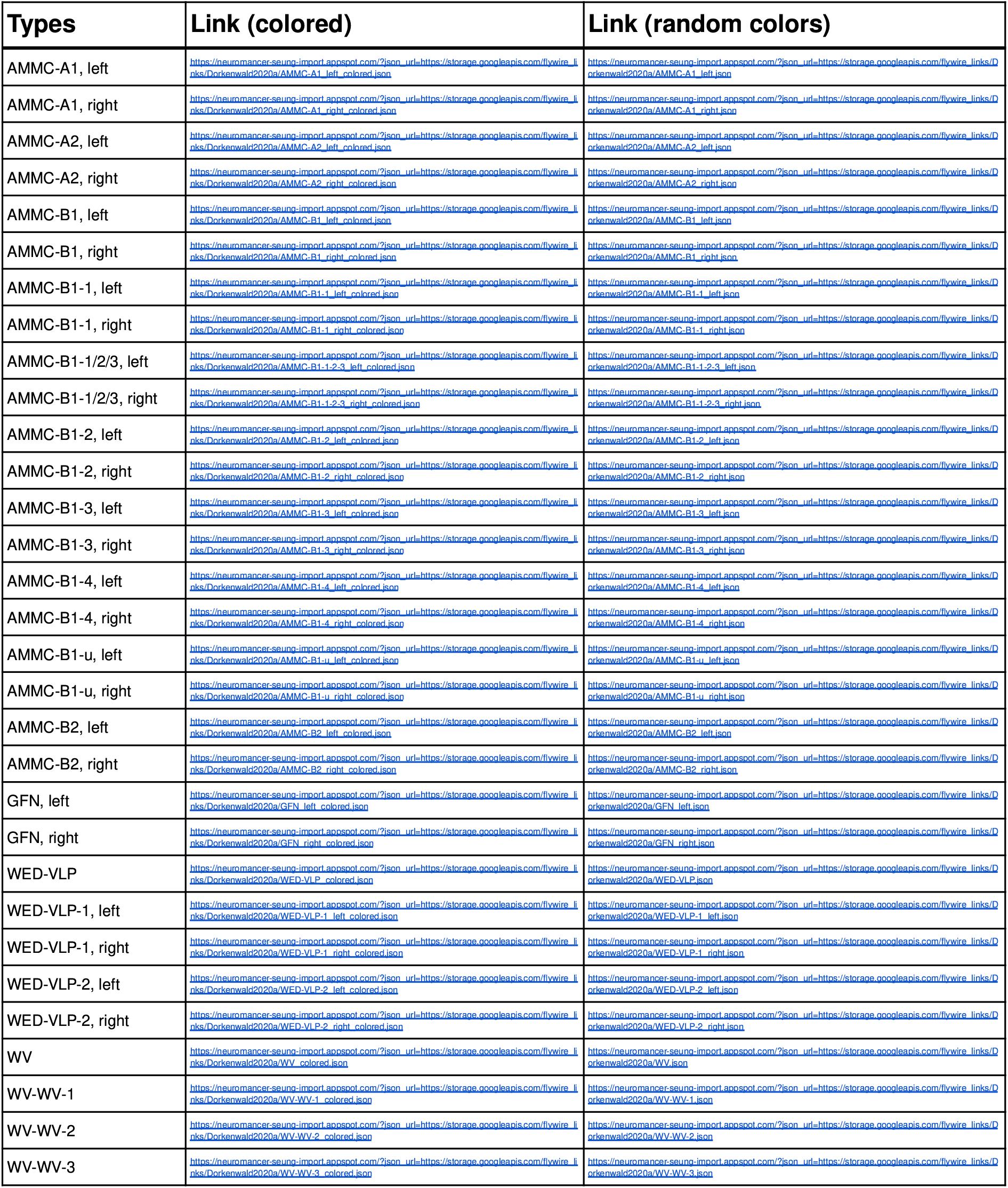
Links to view neurons in Neuroglancer. Links forward users to our frontend and present neurons in 2D and 3D. No login is required to view these neurons; the selection of other neurons is, however, not possible without signing up to FlyWire and onboarding. Links are best viewed in a Chrome browser.

## METHODS

### Alignment

We started from the aligned dataset published by Zheng et al. (Zheng et al. 2018)(v14). Using the method outlined in Mitchell et al. (Mitchell et al. 2019) we trained neural networks through self-supervision to predict pairwise displacement fields between neighboring sections. Here, every location stores a vector pointing to its source location. We introduce a smoothness regularization into the training to ensure continuous transformations. This prior is relaxed at image artifacts such as cracks and folds. We first trained a convolutional neural network to detect image artifacts from a manually labeled training set, then used the predicted masks to adjust the smoothness prior during training of the displacement field network. We combine the pairwise displacement fields to generate a displacement field for every section, and apply the result to the data to create a newly aligned stack.

### Cross alignment registration and brain renderings

Our alignment created a vector field for transformations from FlyWire’s space (v14.1) to the original alignment space (v14). In order to transform data from v14 into v14.1 (e.g. synapses and brain renderings), we created an inverse transformation of the vector field at a resolution of 64 × 64 × 40 nm. Locations in v14 are then transferred to v14.1 by applying the closest displacement vector from the inverse transformation.

The v14 brain rendering was acquired from the hemibrain website: https://flyconnectome.github.io/hemibrainr/reference/hemibrain.surf.html. The v14 whole brain neuropil rendering was acquired from the virtual fly brain website: https://fafb.catmaid.virtualflybrain.org/.

### Segmentation ground truth

We made use of the publicly available ground truth from the CREMI challenge (https://cremi.org) to train our convolutional neural network for predicting affinities. We realigned these ground truth blocks as they contained misalignments as well.

### Segmentation

We applied our segmentation pipeline as outlined in (Dorkenwald et al. 2019; Lee et al. 2017) without the use of long-range affinities. Additionally, we introduced a size-dependent threshold to break big, dumbbell shaped mergers occuring at low threshold. In the affinity graph, we ignored any edges between two large segments *s1*, *s2* if *mean(affinities(s1, s2)) < 0.5* and *min(s1, s2) > 1,000* and *max(s1, s2) > 10,000* representing supervoxel counts.

### Proofreading

The ChunkedGraph is our backend database structure that enables parallel proofreading at scale. It stores the mean affinity graph of supervoxels and uses an octree to provide a spatial embedding for fast updates of the connected components representing neuronal segments. We adapted the Neuroglancer Frontend to command split and merge operations to our server backend. Every edit is stored with a timestamp allowing earlier states of the segmentation to be displayed at later times. To allow parallel proofreading of multiple users, we implement per-neuron locks where we process edits to the same neuron serially. Our server immediately starts remeshing of changed segments to give the user 3D feedback. Remeshing takes seconds to tens of seconds.

### ChunkedGraph performance analysis

During the beta phase of FlyWire, we measured server response times for various requests by all users (Fig. 3e,f). These numbers reflect real interactions and are affected by server and database load and are therefore an underestimate of the capability of our system.

We used real split edits as the basis for the comparison of the ChunkedGraph with a naive implementation that had been performed in FlyWire’s beta phase prior to this analysis. For this comparison, we used the same BigTable table but ignored the additional ChunkedGraph hierarchy for the naive implementation.

### Proofreading evaluation

183 neurons were proofread by 13 proofreaders consisting of both scientists and expert tracers from the Seung and Murthy labs. For 49 of these we measured the time it took the neuron in the first round. For all 183, we only proofread major components (backbones) and did not attempt to look for orphan twigs, small microtubule-free terminal branches, as defined in (Schneider-Mizell et al. 2016). However, proofreaders still added those that were immediately obvious to them.

To obtain the number of edits for each neuron, we excluded edits made to chop neurons apart for inspection, that were later reversed by merging those pieces back together. We also excluded edits to a segment that was removed from that neuron later in the proofreading process. Consider the example where a neuron was merged to a big component containing other neuronal segments. We did not count edits for removing other neuronal segments from that component towards the edit count for the neuron at hand. More specifically, we only considered merge operations where all merge locations remained in the neuron at the end of proofreading and split operations where exactly one side of the split was contained in the final neuron.

We calculated the volumetric change of edits and the volumetric completeness from the segmentation by collecting the supervoxels that were added/removed and adding up the voxel within each of them. We then multiplied this number with the nominal resolution of the segmentation (16 × 16 × 40 nm).

### Twig and backbone synapse evaluation

We randomly collected 999 synapses from all synapses predicted by Buhmann et al. (Buhmann et al. 2019). One expert evaluated all synapses as true positive (615), false positive synapses (285) or ambiguous (99). Next, this expert evaluated the reconstructions of the pre- and postsynaptic sides of the true positive synapses as either belonging to a twig that was attached to a backbone (“twig - attached”), twig that was not attached to a backbone (“twig - orphan”) or “backbone.”

### Identifying all cells within a class

We aimed to find every cell of the mechanosensory types investigated here. To do so, a location was chosen in the soma tract of a cell lineage, where proximal neurites were tightly grouped into a clear bundle, often surrounded by glia. Alternatively, in some cell types without tightly clustered proximal neurites, a location was chosen in a distinctive region of the backbone where these cells showed bundling. By examining in XY, YZ and XZ, a view was chosen that displayed the bundle in cross-section, to ensure that all cells in the bundle were visible. Every neuron in that cross-section was then examined to find the desired cell type. Any neuron that could not be classified was proofread until identification was possible. We expect this approach to reveal most or all cells within a lineage, however there could be reasons why some might be missed (such as a proximal neurite that travels outside the bundle). Locations used: AMMC-A1, right: (103406, 54035, 4640), left: (159748, 56141, 3678). AMMC-B2 commissure: (132104, 71166, 3416). WED-VLP, left: (172786, 69380, 2254), right: (88476, 65205, 3043). WED-WED and AMMC-AMMC (same midline soma tract): (132008, 84118, 4272). AMMC-B1: not all were tightly bundled, used two locations per hemisphere for cross-sections: left: (151298, 69205, 1686) and (152447, 61490, 3218), right: (111828, 67177, 2127) and (111214, 60441, 3615). GF and AMMC-A2: only one cell exists per hemisphere (for AMMC-A2, commissure was examined for confirmation).

### Synapse proofreading

We used automatically detected synapses by Buhmann et al. (Buhmann et al. 2019) for the analysis of the mechanosensory connectome (Fig. 6). During analysis, we noticed a higher occurrence of false positive synapses between some cell types. These were usually cell types that had a high number of contacts due to spatial proximity. We randomly inspected about 25 synapses per <cell type> to <cell type> (eg. AMMC-B1 to WED-VLP or AMMC-B1 to AMMC-B1) and disregarded connections with mostly false positive or questionable synapses. This exclusion mostly affected connections within cell types (eg. AMMC-B1 to AMMC-B1). We did not remove single false positive synapses and the remaining <cell type> to <cell type> connections reported in Figure 6 still have false positive synapses among them.

### Celltype division

We divided cell-types into sub-types according to their connectivity and then verified the subdivision morphologically. We list all numbers in Supplementary Table 1.

#### WED-VLP

Neurons receiving more than 10 synapses from the ipsilateral AMMC-A2 were classified as WED-VLP-1, all others as WED-VLP-2.

#### AMMC-B1

We first selected neurons with more than 50 synapses onto any WED-VLP. These were then labeled as AMMC-B1-1 if they made more than 50% of their WED-VLP synapses onto WED-VLP-1 and AMMC-B1-2 otherwise. Out of the remaining AMMC-B1 neurons (not −1 or −2), those with more than 80 synapses onto any WV-WV neuron were labeled as AMMC-B1-3. From the remaining AMMC-B1 cells, we labeled those as AMMC-B1-4 if they made at least 20 synapses onto AMMC-A1, AMMC-A2 and GFN cells combined. The remaining cells were classified as AMMC-B1-u.

#### WV-WV

First, we labeled all WV-WV neurons with more than 20 synapses onto AMMC-A1, AMMC-A2 and GFN combined as WV-WV-3. Out of the remaining neurons, we labeled those with more than 100 synapses onto WED-VLP as WV-WV-1. WV-WV-2 was made up of all remaining WV-WV neurons.

## ACKNOWLEDGMENTS

HSS and MM acknowledge support from NIH BRAIN Initiative RF1 MH117815-01 to HSS and MM. HSS further acknowledges support from the Mathers Foundation, as well as assistance from Google and Amazon.

We are grateful for support with FAFB imagery by Stephan Saalfeld, Eric Trautman, and Davi Bock. We thank Greg Jefferis, Davi Bock, and Albert Cardona for advice regarding the community. We thank Greg Jefferis and Philipp Schlegel with help with the brain renderings. We thank Garrett McGrath for computer system administration, and May Husseini for project administration. We are grateful to J. Maitin-Shepard for neuroglancer. We are grateful to J. Buhmann and J. Funke for discussions and for making their automatically predicted synapses publicly available. We thank Nuno da Costa and Agnes Bodor for feedback on the proofreading system during its development. We thank the Allen Institute for Brain Science founder, Paul G. Allen, for his vision, encouragement and support.

This work was supported by the Intelligence Advanced Research Projects Activity (IARPA) via Department of Interior/ Interior Business Center (DoI/IBC) contract numbers D16PC00003, D16PC00004, and D16PC0005. The U.S. Government is authorized to reproduce and distribute reprints for Governmental purposes notwithstanding any copyright annotation thereon.

Disclaimer: The views and conclusions contained herein are those of the authors and should not be interpreted as necessarily representing the official policies or endorsements, either expressed or implied, of IARPA, DoI/IBC, or the U.S. Government.

## CONTRIBUTIONS

TM, NK realigned the dataset with methods developed by EM, BN, TM and infrastructure by SP, ZJ, JAB, SM wrote code for masking defects and misalignments. KL trained the convolutional net for boundary detection, using ground truth data realigned by DI. JW used the convolutional net to generate an affinity map which was segmented by RL. SD, NK created the proofreading system with input from JZ and ZA. NK, MAC, OO and AH adapted and improved Neuroglancer for proofreading and annotations. SD, FC, CSM, CSJ, DB built the server infrastructure to host FlyWire and manage users. WMS ingested the images into cloud storage. CM managed the community, designed the training tutorials and trained proofreaders. CB, JG, DD, LER, SK, AB, JH, MM, SM, BS, KW, RW tested the site and proofread neurons. CM, JG devised neuron annotation procedures. SCY managed proofreaders and evaluated twigs. SD evaluated the proofreading system. SD, CM analyzed the data. SD, CM, HSS, MM wrote the manuscript. HSS, MM led the effort.

